# Efficient biological design via genomic in-context learning

**DOI:** 10.64898/2026.10.28.762643

**Authors:** Garyk Brixi, Yukun A. Hao, Divya Nori, Aditi Merchant, Talal Widatalla, Samuel H. King, Katie E. Shalvarjian, Dan M. Park, Brian L. Hie

## Abstract

In-context learning (ICL), the ability to perform new tasks from a few examples supplied in a prompt, is a defining emergent capability of large language models. Here, we show that ICL emerges in the genome language model Evo 2 and enables biological design. With as few as four examples in the prompt and no additional training, Evo 2 generates realistic metal-binding proteins, *de novo* antibodies, protein-RNA systems, and entire bacteriophage genomes. For ΦX174-like bacteriophage design, ICL surpasses previously fine-tuned models on computational measures of design quality and diversity. Beyond ΦX174, conditioning the model with different phage prompts enables us to efficiently design MS2 and M13 genomes *in silico* without fine-tuning. Iteratively updating the prompt with newly generated phage genomes, which we refer to as genomic chain-of-thought, further improves the diversity of designs while maintaining quality metrics. Paired with an inference engine that accelerates generation up to 35-fold and extends practicable context to a megabase, these results establish ICL as a versatile and efficient paradigm for biological sequence design.

## Introduction

Genome language models learn from DNA sequences shaped by billions of years of evolution and can predict properties of DNA, RNA, or proteins from individual genes to entire genomes (Brixi et al., 2026; Ye et al., 2026; King et al., 2026; Li et al., 2026; Cornman et al., 2024). Yet directing these models to generate sequences with desired functions remains challenging. Existing approaches often rely on fine-tuning with thousands of task-specific examples (King et al., 2026) or on functional associations encoded in genomic neighborhoods (semantic design; Merchant et al., 2026). Large language models (LLMs) trained on next token prediction exhibit emergent capabilities that were never explicitly trained for, such as in-context learning (ICL), in which few-shot prompted examples can improve the model response to diverse new tasks (Brown et al., 2020). Due to its data efficiency and simplicity, ICL allows flexible steering of model generation, underlying chain-of-thought prompting (Wei et al., 2022; Kojima et al., 2022) and reasoning in LLMs (OpenAI et al., 2024; DeepSeek-AI et al., 2025).

Evo 2 exhibits ICL on synthetic symbolic tasks (Breslow et al., 2025), but the broader potential of this capability for biological design remains largely unexplored. In a companion study, we designed and experimentally validated mammalian regulatory elements using ICL (Hao et al., 2026). We therefore hypothesized that providing example sequences in the prompt could teach genomic models general principles of evolutionary diversification, such as mutation and recombination, improving biological design across scales.

Here, we use genomic ICL to design lanmodulins, rubisco, anti-CRISPR proteins, epitope-specific *de novo* antibodies, CRISPR–Cas systems, and mitochondrial and bacteriophage genomes. To make this approach practical across sequence lengths, we developed an efficient inference engine for Evo 2. Across the evaluated tasks, we achieved comparable or better *in silico* pass rates on generating novel and coherent sequences compared to baselines generated by task-specific, fine-tuned models or by semantic design. By recursively prompting Evo 2 with computationally evaluated generated sequences, we observed that ICL enables improvement of design quality and diversity, which we refer to as a genomic chain of thought (CoT). Together, genomic ICL and CoT enable complex biological design from only a handful of example sequences, establishing a promising new paradigm for generative biology.

## Results

### Efficient in-context learning with accelerated Evo 2 inference

In a parallel paper, we have shown that Evo 2 learns to generate functional regulatory DNA sequences by simply concatenating in-context examples in the model’s prompt (Hao et al., 2026). We further hypothesized that ICL can offer a data- and compute-efficient way for Evo 2 to generate proteins, protein-RNA systems, and even whole genomes. Testing this hypothesis across diverse biological systems requires efficient model inference (**Fig. 1A**). However, the published inference engine for Evo 2, vortex, was not optimized for such applications and is resource demanding. Open-source inference engines, such as vLLM and SGLang (Kwon et al., 2023; Zheng et al., 2023), offer a memory- and compute-efficient way to serve attention-based natural language models, but lack direct support for genome-customized hybrid models such as Evo 2. To overcome this challenge, we adapted Evo 2’s architecture for vLLM compatibility and further accelerated inference with kernel optimization (Hao, 2026) and speculative decoding (Li et al., 2024).

**Fig. 1.**
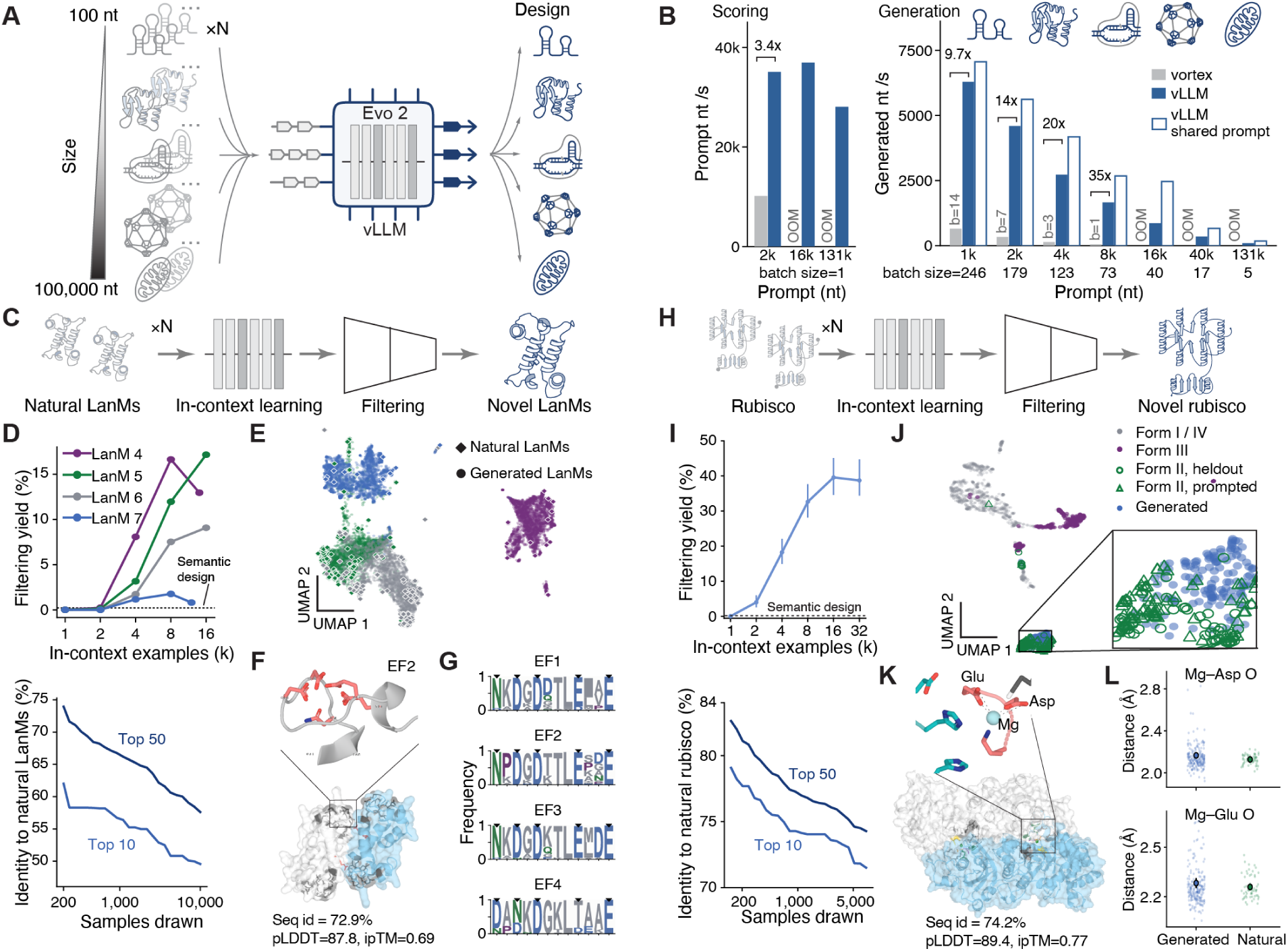
Efficient in-context learning of natural proteins. **(A)** In-context learning allows Evo 2 to design novel biological systems from a few examples, from ∼100 nucleotides to ∼100,000 nucleotides. **(B)** Optimizing Evo 2 with the vLLM engine increased the speed of both prefilling (scoring) and decoding (generation). Speed was obtained on an H100 GPU (80GB). OOM, out-of-memory. For decoding, we used the largest batch size allowed by the GPU. **(C)** In-context learning of lanmodulins (LanMs). **(D)** Test-time scaling of Evo 2 in designing novel LanMs. Top: Percentage of sequences passing a set of filters for HMM search, sequence novelty, and sequence diversity as a function of number of in-context examples. We prompted Evo 2 with CDS of natural LanM from each different functional cluster. Below: Averaged top sequence novelty scales as more samples are drawn. **(E)**UMAP of ESMC embedding of generated LanMs and natural LanMs. Colors are the same as the upper panel of (D). Diamonds indicate natural LanMs and round dots indicate generated LanMs. **(F)**Predicted dimeric structure of a designed LanM4 with sequence identity of 72.9% to the closest natural LanM. As a reference, Hans-LanM has a predicted ipTM of 0.67. The enlarged structure shows the five residues on the second EF hand that may coordinate with lanthanide ions. **(G)** Frequency of residues on four EF hands in generated LanM4 proteins. Black arrows indicate the conserved sites in the Hans-LanM cluster. **(H)** In-context learning of rubiscos. **(I)** Test-time scaling of Evo 2 in designing novel rubiscos. Panels similar to (D). Error bar: 95% confidence interval. **(J)** UMAP of ESMC embedding of generated and natural rubiscos. **(K)** Predicted dimeric structure of a designed rubisco with sequence identity of 74.2% to the closest natural rubisco. Enlarged structure shows that Asp and Glu coordinate with an Mg^2+^ ion. **(L)** Distribution of distances between the Mg^2+^ ion and the two coordinated residues Asp and Glu. Distribution in generated rubiscos is similar to distribution of natural rubiscos. Each dot is a generated or natural rubisco.

The resulting inference engine (Evo2-flash) with converted Evo 2 models exhibits significant gains in both inference speed and memory efficiency. Evo 2 7B generates sequences with up to 7,000 bases per second on an H100 GPU, accelerating generation from 9.7× to 35× compared to Evo 2 with vortex and easing the constraint on GPU memory, depending on the batch size and context length (**Fig. 1B**). For 8,192-nt to 16,384-nt context length, sufficient for ICL of proteins, Evo 2 samples 1600 to 4000 bases per second with normal decoding (**Fig. 1B**). At medium context, such as 40,960-nt context length, sufficient for ICL of phage genomes, Evo 2 samples 350 to 664 bases per second whereas vortex raises out-of-memory (OOM) errors (**Fig. 1B**). For long contexts at 131,072 nt or higher, sufficient for ICL of mitochondrial genomes, Evo 2 samples 95 to 177 bases per second (**Fig. 1B**). A trained drafter for speculative decoding further improved generation speed for long contexts (**fig. S1A**). Moreover, Evo2-flash supports a wider range of GPUs, such as the RTX 3090 and the L4 and A100 GPUs available on Google Colab, and extended prompt length to 1 megabase on a single H100 GPU (**fig. S1B**). These optimizations make genomic ICL more accessible and practical across scales.

### Generative in-context learning of natural proteins and protein-RNA systems

We first sought to test the genomic in-context-learning hypothesis in natural proteins—using lanmodulin (LanM) (Mattocks et al., 2023; Diep et al., 2026), ribulose-1,5-bisphosphate carboxylase (rubisco) (Bar-On and Milo, 2019), and anti-CRISPR (Acr) proteins (Davidson et al., 2020; Merchant et al., 2026) as testbeds— by prompting Evo 2 with concatenated coding sequences (CDSs) of natural proteins separated by a special nucleotide sequence.

Lanmodulins (LanMs) are a family of bacterial proteins that selectively bind lanthanide ions (Cotruvo et al., 2018). High-throughput characterization of natural LanMs has identified eight functional clusters distinguished by their relative selectivity across the lanthanide series (Diep et al., 2026). Among these, Cluster 4 is particularly promising for rare-earth element separation, containing members such as *Hansschlegelia quercus* LanM (Hans-LanM) (Mattocks et al., 2023) and *Ancylobacter lacus* LanM (Al-LanM) (Choi et al., 2026), which can discriminate between chemically similar lanthanide ions. However, Cluster 4 itself contains only 14 proteins, offering a limited repertoire of natural sequences for further development (Diep et al., 2026). To expand this repertoire, we prompted Evo 2 with coding sequences from LanMs from four of the eight functional clusters (Clusters 4–7). We screened the generated sequences using profile hidden Markov models (HMMs) to identify LanM-like proteins with plausible EF-hand architectures, then applied additional filters for novelty relative to natural LanMs (<80% protein sequence identity) and diversity among the designs (**Fig. 1C**). Across the conditions tested, up to 15% of sequences generated with Cluster 4 prompts passed all filters, with an overall throughput of >6,000 sequences passing all three filters per GPU-hour (**Fig. 1D**). Increasing the number of prompted examples improved the yield of designs passing the filters, while increasing the sampling budget enabled the discovery of progressively more novel sequences. Specifically, sequence identity to natural LanMs among the 50 most novel designs decreased by approximately 3.8% with each doubling of the number of sampled sequences, demonstrating test-time scaling of sequence novelty in ICL-based protein design (**Fig. 1D**).

The generated proteins also exhibited features associated with their target clusters. Dimensionality reduction of embeddings from the ESMC protein language model placed generated sequences near natural LanMs from the corresponding functional clusters (**Fig. 1E**; Candido et al., 2026). ESMFold2 models of a Cluster 4 design exhibited homodimeric architectures, consistent with the experimentally observed dimerization of Hans-LanM and Al-LanM (Mattocks et al., 2023; Choi et al., 2026) (**Fig. 1F**; **fig. S2A**). These designs further recapitulated the characteristic residue patterns across all four EF-hand motifs of natural Cluster 4 LanMs (**Fig. 1G**). Together, these observations are consistent with Evo 2 capturing cluster-associated sequence features from in-context examples while generating candidates distinct from known natural LanMs.

We next extended genomic ICL to form II rubisco, prompting Evo 2 with coding sequences of proteins longer than 400 amino acids (Kehl et al., 2026; Jaffe et al., 2019). We applied a similar set of filters to extract novel rubisco-like sequences (**Fig. 1H**), and obtained ∼1900 designs per GPU hour at a protein sequence identity threshold of less than 85%. Genomic ICL achieved a higher proportion of designs passing all filters compared to semantic design, in which Evo 2 was prompted with the genomic contexts of rubisco genes. This yield increased with the number of examples provided in the prompt (**Fig. 1I**). Increasing the sampling budget also enabled the discovery of progressively more divergent sequences, extending test-time scaling of sequence novelty to a second protein family (**Fig. 1I**). UMAP visualization of ESMC embeddings placed the generated sequences alongside natural form II rubiscos, including sequences held-out from the prompts (**Fig. 1J**). The structural models of one representative design exhibited homodimeric architectures with plausible Mg^2+^ coordination (**Fig. 1K**; **fig. S2B**). Further analysis of the predicted functional sites revealed Mg^2+^ coordination geometries resembling those of natural rubiscos (**Fig. 1L**). These results extend genomic ICL to enzyme design, yielding novel sequences that recapitulate key sequence features and predicted structural characteristics of natural form II rubiscos.

We next asked whether Evo 2 ICL could generate members of highly diverse protein families. We selected anti-CRISPR (Acr) proteins, which inhibit CRISPR systems through diverse mechanisms and exhibit extensive sequence variation. Although previous work showed that Evo 1.5 could generate functional Acrs through semantic design using their surrounding genomic context (Merchant et al., 2026), it remained unclear whether diverse Acr sequences alone could provide effective conditioning through ICL. We therefore supplied coding sequences from either type II Acrs or the broader set of types I–VI and evaluated generation success as the number of examples increased. Consistent with greater sequence similarity within a single type, candidate yield improved over semantic design baselines with 32 examples for type II Acrs, whereas enrichment across types I–VI became apparent only at 128 examples (**Fig. 2A**). At this example number, 47% and 23% of generations, respectively, passed computational Acr filters, compared with 6% for semantic design, indicating that sufficiently large conditioning could support ICL from highly diverse examples. The broader type I–VI set also yielded Acr candidates with a wider distribution of sequence identities to known proteins (**Fig. 2B**), including candidates with only remote similarity to known Acrs (**Fig. 2C**), although divergence remained lower overall than with semantic design. To examine the effect of prompt diversity more directly, we grouped examples from a single Acr family, AcrIIA9, by their median pairwise sequence identity and fixed the number of prompt examples at 64. Consistent with the broader Acr ICL results, greater variation among examples generally increased divergence of candidate Acrs from the prompt but lowered the fraction of generations passing Acr filters (**Fig. 2D**). Together, these results demonstrate that genomic ICL can generate diverse candidates from highly variable protein families, with example selection offering a means to tune candidate yield and sequence novelty.

**Fig. 2.**
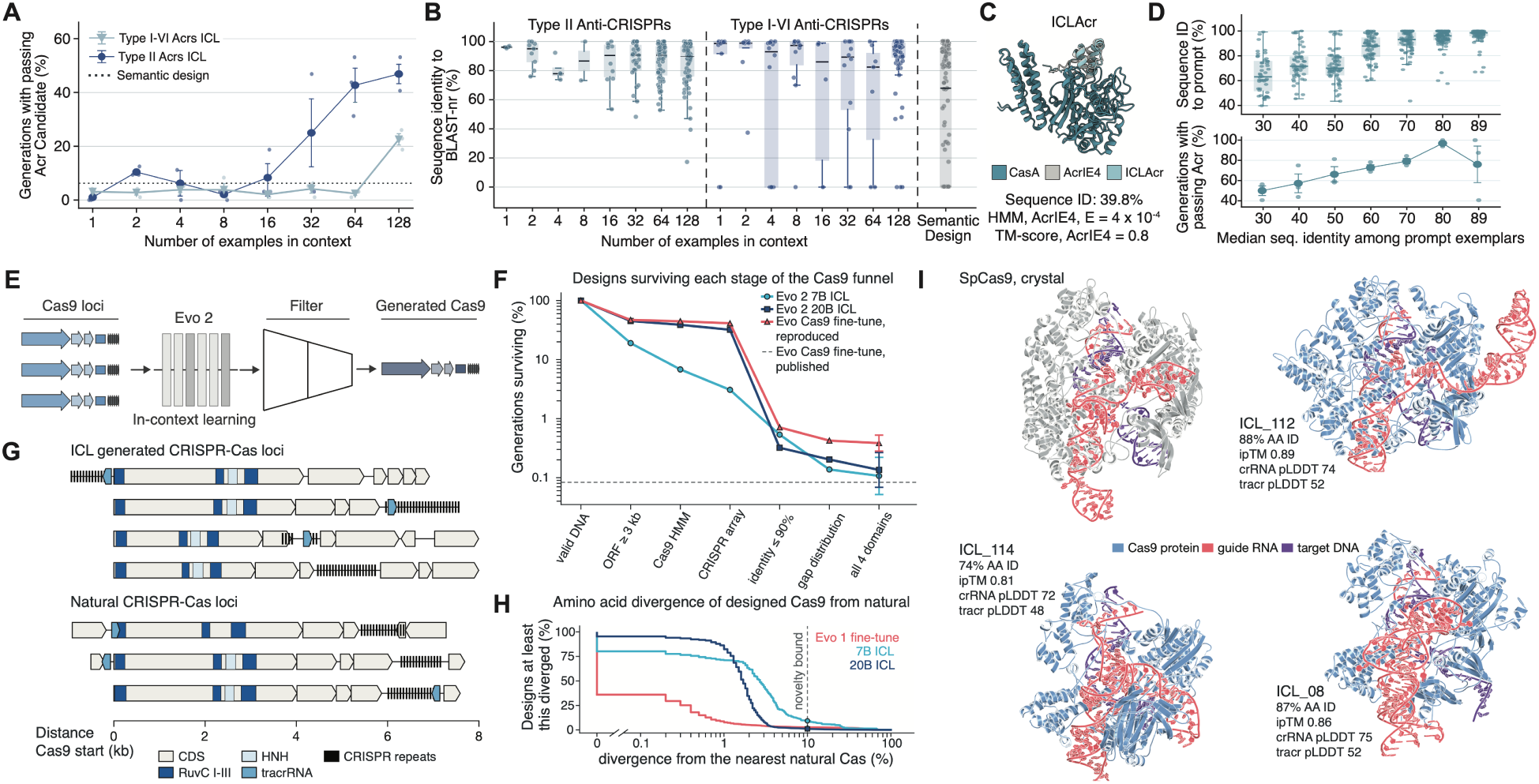
In-context learning of highly diverse proteins and protein-RNA complexes. **(A)** Filter passing rate of generated Acrs as a function of number of prompted examples. **(B)** Sequence novelty of generated Acrs as a function of number of prompted examples. **(C)** Predicted structure of a generated Acr with 40% sequence identity to closest known Acr in complex with CasA. **(D)** Sequence novelty and passing rate as a function of sequence diversity of prompted AcrIIA9 examples. **(E)** In-context learning of CRISPR-Cas operons. **(F)** Pass rate after each stage of the filters across several methods. **(G)** Generated CRISPR-Cas loci resemble natural loci. **(H)** Percent of designed Cas9 as a function of amino acid sequence divergence from the nearest natural Cas. **(I)** AlphaFold 3 predicted structure of ICL-designed Cas9 proteins alongside DNA and guide RNA compared with the SpCas9 crystal.

We next tested whether ICL could generate CRISPR-Cas loci predicted to contain a CRISPR array, transactivating CRISPR RNA (tracrRNA), and Cas9 protein. As neither Evo 1 or Evo 2 were able to generate Cas9 when prompted with partial loci, generation of EvoCas9 required fine-tuning Evo 1 on tens of thousands of natural Cas9 loci before sampling and filtering thousands of candidates (Nguyen et al., 2024). Here, we instead supplied just four natural Cas9 loci in-context as examples, generated designs, and evaluated them using previously published filtering criteria for correctness and sequence diversity (Nguyen et al., 2024).

ICL for CRISPR-Cas yielded a lower per-sample pass rate compared to fine-tuning (0.14% versus 0.39%, **Fig. 2F**). However, because fine-tuning Evo 1 consumed 881 H100 hours, producing a library of 64 passing designs cost roughly 900 H100 hours compared with 81 hours using Evo 2 7B ICL. Thus, despite the lower yield, ICL enables an order-of-magnitude reduction in time. The generated loci contain expected elements and architecture, including the CRISPR array, tracrRNA, and Cas9 protein (**Fig. 2G**). ICL-generated Cas9 genes are more than 10% distant from their nearest natural homolog 9.3% of the time compared with 2.7% of finetuned Evo 1 generations (**Fig. 2H**). AlphaFold 3 predicts the generated Cas9 proteins to adopt folds spanning multiple natural variants (**Fig. 2I**).

We further extended genomic ICL to entire organelle genomes by prompting Evo 2 with human, primate, or mammalian mitochondrial genomes (**fig. S3A**). By increasing numbers of examples in the prompt, ICL generates sequences that recapitulate the characteristic 37-gene complement of mammalian mitochondrial genomes (**fig. S3B**). The diversity of generated sequences reflected that of the prompt population: human mitochondrial genomes yielded less diverse designs, whereas broader primate or mammalian prompts yielded more diverse designs (**fig. S3, C and D**). These results extend genomic ICL to organelle-scale design, with prompt composition providing a means to modulate sequence diversity.

### Generative in-context learning of epitope-targeting antibodies

Designing antibodies against a specified epitope remains a central challenge in therapeutic protein design. Recent structure-based generative methods, including Germinal, can design functional *de novo* antibodies against diverse sets of antigens (Mille-Fragoso et al., 2026; Pacesa et al., 2025). However, the computational cost of these methods is still substantial. For example, Germinal required a median of 23.5 GPU hours to design 8 hits per target, and many targets require more sampling. Additionally, structure-based generative pipelines typically cannot improve designs using new computationally or experimentally validated hits.

We sought to test if ICL with Evo 2 could rapidly design novel epitope-specific hits by learning in context from 8 *de novo* binders without any explicit structural conditioning. We compared against an antibody-specific structure-conditioned sequence model, AbMPNN (Dreyer et al., 2023); a PSSM baseline using aligned prompt examples’ per-position residue statistics; and continued sampling from Germinal. We assessed methods on predicted binding quality as well as structural diversity of the generated sequences, reflecting the goal of creating a diverse set of binders on a fixed compute budget (**Methods**).

Under this evaluation, ICL produced more distinct filter-passing designs than any other method (**Fig. 3B**), and was the only method to yield at least one hit for every antigen tested. We also observed greater structural diversity of designs compared to structure-conditioned sequence generation with AbMPNN, though this diversity was less than that observed with Germinal when sampling the same number of sequences (**Fig. 3C**). We note that with Evo2-flash, we can generate over 13,000 designs per GPU-hour, matching the throughput of AbMPNN and shifting the bottleneck in a computational design campaign from sequence generation to structure prediction (**Fig. 3D**). When designs were required to be as structurally distinct from one another as two independently sampled Germinal designs, ICL retained the most passing designs. Inspecting individual designs showed that ICL composes new sequences by recombining segments of in-context examples and introducing novel mutations (**Fig. 3E**), resulting in designs with high predicted interface confidence scores while being novel at the CDR sequence level (**Fig. 3F**). Notably, hit classification requires the design to contact the desired epitope, suggesting Evo 2 can produce epitope-targeted antibodies from prompted examples alone, without any explicit structural information.

**Fig. 3.**
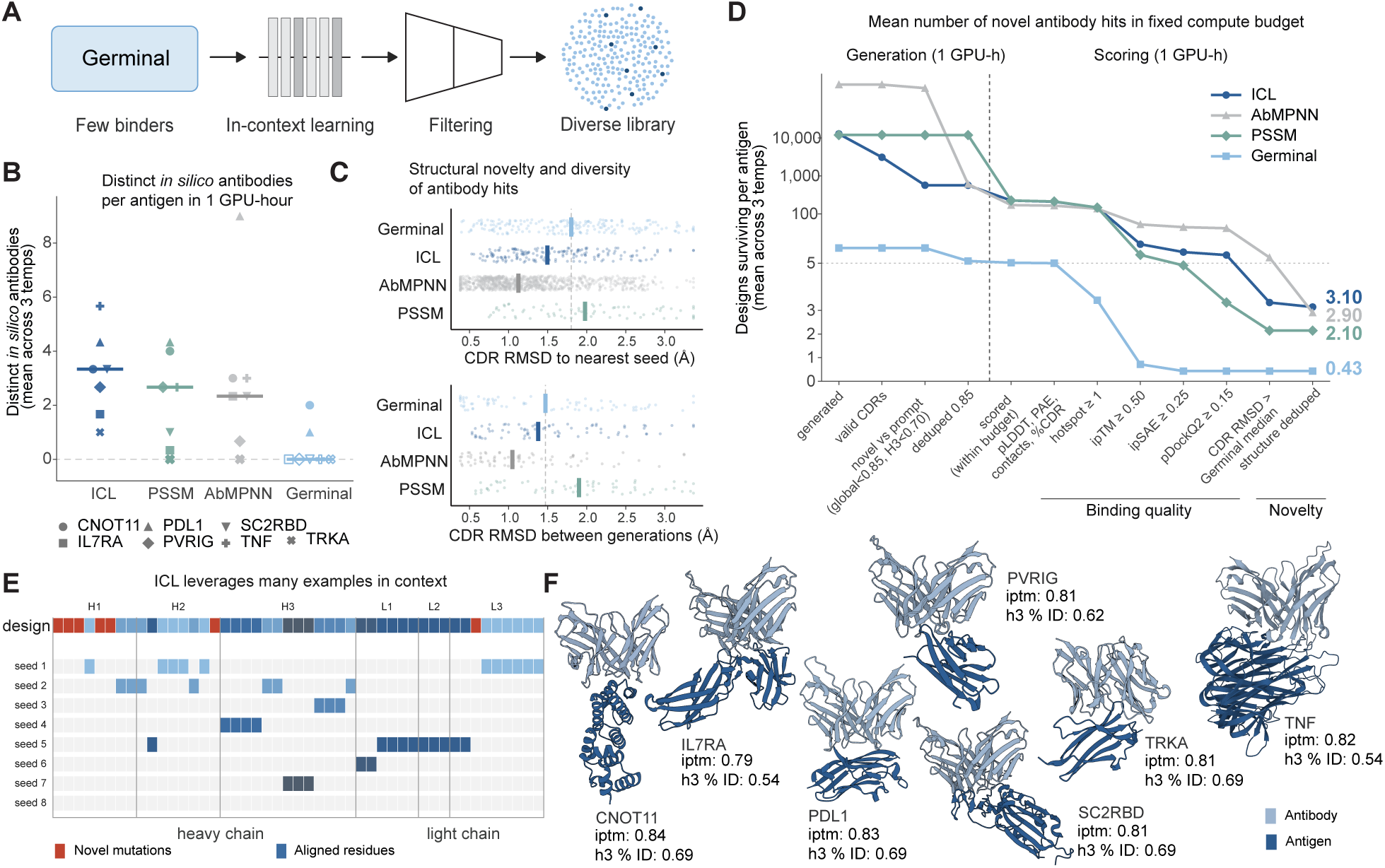
In-context learning of antibodies. **(A)** Schematic showing how sequences generated with Germinal are an input to Evo 2 ICL and then filtered to create a diverse library of designs. **(B)** The number of distinct *in silico* passing binders generated in 1 GPU hour per antigen, showing heterogeneity between antigens compared between ICL, PSSM, AbMPNN, and Germinal. CNOT11, CCR4–NOT transcription complex subunit 11. IL7RA, Interleukin-7 receptor subunit alpha. PDL1, Programmed death-ligand 1. PVRIG, Poliovirus receptor-related immunoglobulin domain-containing protein. SC2RBD, SARS-CoV-2 spike protein receptor-binding domain. TRKA, Tropomyosin-related kinase A. TNF, Tumor necrosis factor. **(C)** The structural novelty and diversity based on CDR RMSD to seed and CDR RMSD between different generated sequences compared between the four design methods. **(D)** The number of designs passing each filtering step when using the different design methods with a one hour generation budget and a one hour scoring budget. **(E)** Example generated design showing how ICL recombines the different examples in context and introduced new variation. **(F)** Predicted AlphaFold structures of seven design targets and Evo 2 generated antibodies. Abbreviations for design targets are the same as (B).

### Whole-genome design with in-context-learning and genomic chain-of-thought

Designing entire bacteriophage genomes requires generating DNA, RNA, and proteins which give rise to organismal function. Evo 2 previously required extensive fine-tuning on a large training set to generate ΦX174-like phages (King et al., 2026). We reasoned that Evo 2 may instead design realistic phages through in-context learning, conditioning only on four or eight natural genomes placed in the prompt, without weight updates (**Fig. 4A**). We evaluated the sequences *in silico* using the correctness and novelty metrics established for generated ΦX174-like phages (**Methods**; King et al., 2026). Quality filters assess if the phage is plausible, while novelty filters measure divergence from natural *Microviridae* and all phage proteins.

**Fig. 4.**
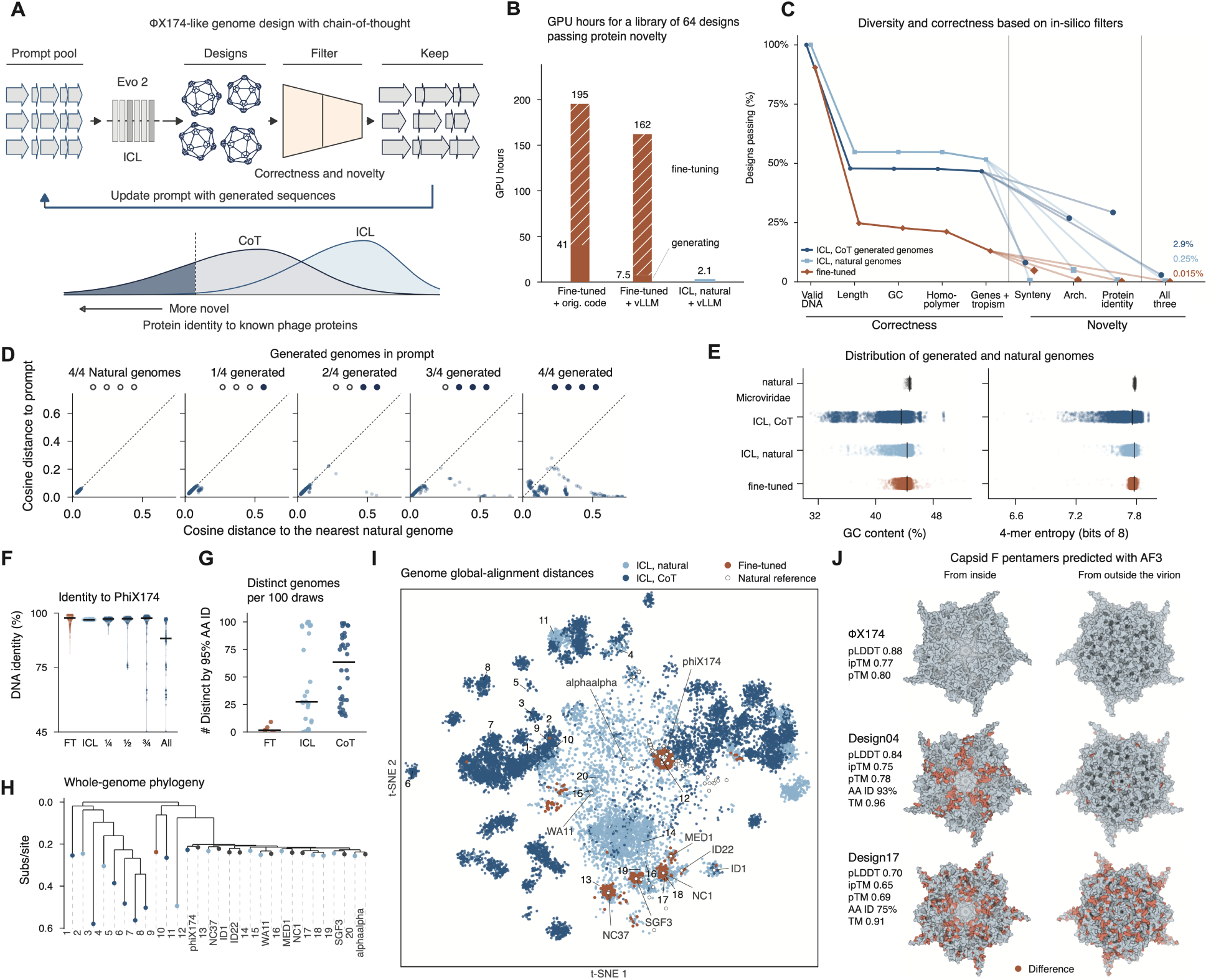
Genomic chain-of-thought design of bacteriophage genomes. **(A)** Schematic showing in-context learning and chain-of-thought for whole-genome generation. **(B)** Estimated GPU time to generate a library of 64 ΦX174-like phage genomes which pass the *in silico* correctness filters, a nucleotide identity of ≤ 98.9% to natural genomes, and a mean protein identity cutoff of ≤ 95% to best-matching PHROG database hits. **(C)**The percentage of ΦX174-like generations by strategy that pass each correctness and novelty filter compared between Evo 2 fine-tuned, Evo 2 ICL, and Evo 2 CoT. **(D)** Sequence diversity as CoT proceeds and generated genomes replace natural examples in a four-genome prompt (first-in, first-out). Each point represents a correctness passing genome, with 6-mer cosine distance to prompt and to the nearest natural genome. **(E)** Distribution of GC content and 4-mer entropy between natural genomes and those generated from fine-tuning, ICL, or ICL with CoT. **(F)** Distribution of DNA identity to ΦX174 of correctness-passing generated genomes from fine-tuning, ICL, and ICL CoT with progressively more synthetic genomes in the prompt. **(G)** The number of distinct genome clusters (≤ 95% amino acid identity across all genes) generated by 100 correctness passing generations from fine-tuned, ICL, and CoT genomes. **(H)** Whole-genome phylogeny tree with natural genomes

ICL produces correct and diverse phage genomes at a small fraction of the computational cost of using a fine-tuned model. Designing a library of 64 designs that pass all correctness filters and the protein novelty placed alongside generated genomes sampled from the full set of correctness passing designs. **(I)** t-SNE plot based on all-by-all whole-genome global-alignment distances between natural genomes and genomes generated by fine-tuning, ICL, or ICL with CoT. **(J)** ΦX174-like capsid pentamers sampled from generated designs compared with the natural ΦX174, predicted by AlphaFold 3. filter required approximately 200 GPU hours to fine-tune and run inference (162 hours when using the evo2-flash engine), compared with 2.1 GPU hours for ICL on natural genomes (**Fig. 4B**). ICL passed the correctness filters at 51.7% compared with 13.0% for fine-tuning. Moreover, 0.25% of ICL designs pass correctness and all three novelty filters compared with 0.015% for fine-tuning (**Fig. 4C**). The relatively low yield on novelty filters motivates us to further improve ICL.

We hypothesized that iteratively incorporating generated sequences into subsequent prompts, an approach we term genomic chain-of-thought (CoT), could expand design diversity. Genome language models are pre-trained on natural DNA sequences without any reasoning traces, but ICL provides a natural route for conditioning recursively with generated sequences. After sampling a large batch, we append generated genomes that pass filters to the right of the prompt and evict the genomes from the left (first-in, first-out; **Fig. 4A**). CoT therefore differs from ICL in which genomes make up the prompt, and has the same compute cost when prompts are refreshed at the same frequency. As a larger fraction of the prompt are synthetic sequences, the cosine distance both to natural sequences and to the prompt increases (**Fig. 4D**). CoT increased the pass rate for all three novelty filters to 2.9% from 0.25% for ICL, while passing the quality filters at a modestly lower rate to ICL (CoT: 46.7%; ICL: 51.7%). CoT-generated genomes span a far wider range of GC content and 4-mer entropy than natural genomes (GC interquartile range 2.22% vs 0.21%; 4-mer entropy 0.130 vs 0.007 bits), whereas ICL and fine-tuned genomes remain close to natural on both measures (GC 0.41% and 0.37%; entropy 0.010 and 0.019 bits) (**Fig. 4E**).

CoT improves diversity across different metrics while passing quality filters. Measuring the whole genome sequence identity of passing designs against ΦX174, generated genomes are increasingly diverse as more synthetic sequences are used in the CoT (**Fig. 4F**). Measuring the number of distinct genomes based on clusters of 95% amino acid identity, we observe that most sequences form just a few clusters for fine-tuning, while almost half of CoT sequences form different clusters (**Fig. 4G**). As a result of this diversity, the generated genomes can span the entire phylogeny based on whole-genome alignments (**Fig. 4H**). We visualize the global distances between genomes using a t-SNE. While fine-tuning appears to generate many genomes clustered around natural reference genomes, ICL and CoT generate genomes across different sequence neighborhoods (**Fig. 4I**). Despite the diversity, AlphaFold 3 predicted capsid pentamer structures are similar to natural capsids, with TM-scores greater than 0.9 (Zhang and Skolnick, 2004) (**Fig. 4J**), indicating the generations retain ΦX174-like structure.

Having shown that ICL enables efficient design of ΦX174-like genomes *in silico*, we tested whether it extends to phages with different architectures and lifecycles. MS2 is a positive stranded RNA phage with an icosahedral capsid, while M13 is a filamentous single strand DNA phage; both differ fundamentally from the previously designed ΦX174 in genome type, capsid architecture, and lifecycle (Callanan et al., 2021; Knezevic et al., 2021). We developed correctness filters to the genome architecture of M13 and MS2 and assess novelty based on protein identity. For both M13 and MS2, ICL generates genomes that pass correctness filters (M13: 92.8%, MS2: 57.5%), while directly prompting Evo 2 fails to complete either genome (**fig. S4**). Novelty of generated sequences depend on the diversity of natural phages available. For MS2, no generated genomes pass the protein novelty filter, which means no passing sequences to include for CoT. This likely reflects that MS2 genes are within 99% amino acid identity across different genomes (**fig. S4**). However, for the more diverse M13 genomes, 36.6% of genomes passed protein diversity with ICL which further increased to 49.6% with CoT. Together, these results show that with only a few natural genomes in context and no weight updates, Evo 2 can design genomes that pass *in silico* filters across genetically distinct classes of bacteriophage, including an RNA phage.

## Discussion

We report that in-context learning (ICL) emerges in genome language models, enabling data- and compute-efficient biological sequence design across tasks spanning multiple scales of complexity. Given a population of sequences in the prompt, Evo 2 generates new sequences that combine and mutate features of inputs to generate novel candidates with desired biological features across modalities. Using an optimized Evo 2 inference engine, we demonstrate generative genomic ICL for designing lanmodulin, rubisco, CRISPR-Cas operons, human mitochondrial genomes, *de novo* antibodies, and bacteriophage genomes. *In silico* analysis suggests generated sequences are both novel and coherent, with predicted structures and functional features similar to those of natural sequences.

Across these design tasks, Evo 2 ICL generated high-quality and novel sequences with as few as four in-context examples, demonstrating an efficient way to design rare proteins like Hans-LanM. Evo 2 ICL also achieved similar performance as previous fine-tuning campaigns in designing novel CRISPR-Cas operons and bacterial phage genomes without any additional training, which substantially decreases the compute needed for many biological sequence design tasks. Our results suggest that ICL combined with filtering can drive the synthetic evolution of sequences to extend sequence diversity and properties. Feeding generated sequences back into the model as a form of chain-of-thought in DNA enables iterative design of increasingly diverse bacteriophage genomes.

We note that *in silico* metrics do not guarantee biological function, and all evaluations presented here are purely computational. In our companion study, ICL-designed enhancers showed cell-type-specific activity in cultured cell lines and in the mouse brain (Hao et al., 2026), suggesting that generative ICL can produce functional sequences. The computational filters used here provide a consistent basis for comparing ICL against other methods and selecting plausible designs.

Recent papers have discussed using evolutionarily related sequences, possibly arranged as a multiple sequence alignment (MSA), as a way to augment the conditioning provided to protein language models (Yang et al., 2025; Jain et al., 2025), which can be viewed as a case of training for in-context learning. ICL in Evo 2 is an emergent phenomenon that results from training only on natural genomic sequences and enables design of broader biological systems without explicit MSAs.

ICL emerges in Evo 2 during pre-training, consistent with the emergence of analogous phenomena when pre-training sequence models on natural language (Wei et al., 2022; Brown et al., 2020). Post-training towards these capabilities could enhance this behavior, enabling design and composition of novel biological systems in low data settings. Coupled with iterative or automated experimentation, data-efficient frameworks like ICL offer a path toward “laboratory-in-the-loop” design of biological systems at scale.

## Methods

### Re-implementing Evo 2 for the vLLM engine and benchmarking

We developed a vLLM (Kwon et al., 2023) implementation of Evo 2 (Brixi et al., 2026) which improves throughput, memory usage, and performs model surgery to make Evo 2 compatible with broader hardware types. Evo 2 is built on StripedHyena 2 (Ku et al., 2025), a hybrid architecture that mixes Hyena convolution layers with a few attention layers, which vLLM does not support directly. To make it compatible, we re-implemented each Hyena layer as a vLLM state-space-style layer, so that vLLM manages its recurrent state in the same paged memory pool it uses for the attention KV cache. As a result, Evo 2 gets vLLM’s serving features including continuous batching, chunked prefill (with Hyena state carried across chunks), prefix caching, CUDA graphs and speculative decoding. We validated the port against the original implementation with the Evo 2 correctness benchmark. In addition, we implemented custom GPU kernels for the Hyena operations vLLM has no kernels for the long convolutions and recurrent-state computation during prefill, and a fused per-token Hyena step and fused normalization during decoding. These kernels produce identical results to the reference computation.

Evo 2 checkpoints, especially the 1B and 40B/20B variants contain a dependency on the FP8 linear from Transformer Engine (NVIDIA, n.d.), and are only compatible with Hopper hardware. To address this, we used Claude Opus 5.5 and Opus 5 to investigate the cause of the numerical instability and perform model surgery to fix the requirement. Claude identified the specific layers responsible for the numerical inconsistency and proposed and implemented a fix. The dependency turns out to be the accumulation for the FP* tensor-core GEMM, which truncates towards zero in groups of 32 products, which shrinks the first blocks input by 1.4% in the 1B model and 6% in the 20B model, relative to exact accumulation. Using the exact accumulation however, or running other data types like bf16, causes the entire models predictions to be near random, because the first hyena layer effectively cancels most of the token embedding it is added to, so a small difference in the input to the layer leads to a large change in the residual stream. Since this happens in the first layer, the input can be replaced by an exact lookup table over the vocabulary, which was computed once on Hopper and stored in the checkpoint, and can be performed instead of a GEMM on other hardware. This was performed for the 1B and 20B model, but not the 7B which had no such dependency. The 20B model further carried some dependency on FP8 in the final Hyena layer, which led to further, smaller, difference in correctness. The specific channels in this layer where the bf16 and Fp8 forward pass were found to diverge were then performed using an emulated Fp8 GEMM, created by Claude, which is used to patch those columns. This enables Evo 2 20B/40B and 1B models to run correctly on other GPU hardware.

To benchmark the accumulated speedup for Evo 2 7B on an H100 GPU, we used prompts of 1,024 to 131,072 nucleotides constructed as distinct, non-overlapping windows of the *Haloferax volcanii* DS2 genome (Hartman et al., 2010). For each prompt we ran vortex (Poli et al., n.d.) and vLLM at batch size 1 and at the largest batch each engine could fit in GPU memory, and sampled 1,000 nucleotides. Prefill speed was taken from a separate run on the same prompts that generated a single token. Decoding throughput was measured as total generated tokens divided by the wall-clock time, and each value is the median of three runs in fresh processes.

### LanM ICL

We used example LanM sequences of cluster 4, 5, 6, 7 from Diep et al (Diep et al., 2026). Full coding sequences of 1-, 2-, 4-, 8-, 16-shot natural LanMs in each cluster were concatenated and separated by a GGGGGGGG sequence. We construct the prompts such that each *k*-shot example set is a subset of the corresponding 2*k*-shot set. Considering the small number of LanMs from cluster 4 (14) and cluster 7 (12), we did not go beyond 16-shot. For each condition we drew 2,500 samples from 100 distinct prompts for each condition at *T* = 0.8, top-*k* = 4 at a speed of more than 50,000 samples/h. These were filtered for avoiding low-complexity (segmasker (Camacho et al., 2009) X-fraction, *k*-mer repeat coverage, amino-acid diversity and under-representation; 16,697 retained), for EF-hand architecture by Pfam PF00036.39 (Mistry et al., 2021) with pyhmmer (Larralde and Zeller, 2023), and then for novelty against the 560 natural LanM sequences with a threshold of 80% sequence similarity (Diep et al., 2026). We then de-duplicated generated sequences with a threshold of 90% sequence similarity.

### Rubisco ICL

We curated a pool of 147 natural Form II proteins by combining a published source (Jaffe et al., 2025) and UniProt (The UniProt Consortium, 2023) search (search string: ((gene:cbbM) OR (cc_miscellaneous:”form II”)) AND (length:[361 TO 500]) AND (fragment:false)) (1,336-proteins total after removing duplications) and filtered with HMM search (Jaffe et al., 2025). 74 proteins were held out and 73 were used for in-context learning. We reverse-translated to DNA under seeded uniform synonymous codon choice and concatenated with a GGGGGGGG separator. We prompted Evo 2 with 1, 2, 4, 8, 16 and 32-shot rubisco sequences (72 distinct prompts per condition and 8 samples per prompt). Each prompt’s *k*-shot example set is a prefix of the corresponding 2*k*-shot prompt. Sampling used temperature 0.8, top-*k* 4, top-*p* 1.0, and we collected 1500 bases.

We translated the generated sequence in the prompted frame from the ATG start codon to the first in-frame stop and applied the following filters: (1) all valid A/T/C/G bases, (2) length within 361–500 amino acids and no homopolymer longer than 10, and (3) HMM best hit to be Form II with *E* ≤ 1 × 10^−5^ and ≥ 90% profile coverage. Novelty was measured by blastp (Camacho et al., 2009) against the 1,336-protein natural panel, retaining only hits covering ≥ 80% of both query and subject, and additionally requiring the aligning natural to be covered to ≥ 90% to prevent truncated proteins from entering the pool of novel designs. We collected designs below 85% maximum identity to the 1,336-protein panel and de-duplicated them by MMseqs (Steinegger and Söding, 2017) clustering at 85% sequence similarity.

### Acr ICL

To evaluate the ability of Evo 2 ICL to generate anti-CRISPRs, we first curated a pool of anti-CRISPR examples comprising known and high quality candidate Anti-CRISPRs from AcrDB (Huang et al., 2021) and homologs identified in NCBI core_nt, GTDB r232 (Parks et al., 2022), and Swiss-Prot (The UniProt Consortium, 2023) spanning Acr types I, II, III, IV, V, and VI. For each experiment, examples were sampled from the relevant pool (type-I-VI Acrs, type-II Acrs, or AcrIIA9) by cycling through randomly ordered families (type I-VI Acrs and type II Acrs) or example sequences (AcrIIA9) and selecting one sequence per family until the desired shot count was reached. Corresponding genomic coding sequences were concatenated with an ‘ATGCATGA’ separator to construct 1- to 128-shot prompts. For each condition, we used three independent example draws with three prompt replicates each and generated 32 samples per prompt using Evo 2 7B at temperature 0.9, top-*k* 4, and top-*p* 1.0, with 1,500 bp generations.

Candidate anti-CRISPRs were identified following methods adapted from Merchant et al. (2026). Briefly, generations were first scanned with Prodigal (Hyatt et al., 2010) in metagenomic mode to identify candidate ORFs. ORFs were retained if they encoded a protein with a sequence length of 50-200 amino acids, greater than 12 distinct amino acids, and a segmasker low-complexity fraction ≤ 0.1 (Cock et al., 2015). Next, proteins were selected for structure prediction if they had either an Acr HMM hit (E-value less than 0.1) or a logistic prescreen score greater than 0.15. The prescreen combined raw AcRanker (Eitzinger et al., 2020) scores and profile-aware AcrNET (Li et al., 2023) percentiles and was calibrated using known Acrs, other defense proteins, and unrelated prokaryotic proteins. Selected proteins were folded with ESMFold2 (Candido et al., 2026) and retained if their monomer pTM was greater than 0.33. Final Acr candidates were accepted if they received an AcrNET score of greater than 0.99 for proteins with a PSI-BLAST profile or 0.9999 without one. Novelty was assessed by BLASTP against the BLAST non-redundant protein database. Predicted structures were further annotated against PDB100, AFDB50, and BFVD v3 using Foldseek and compared with their closest curated Acr using query-normalized TM-score (Kempen et al., 2024).

### Cas9 ICL

We use natural type II CRISPR-Cas9 loci as in-context examples, using the Cas9 sequence and surrounding context, starting at the Cas9 gene. We restrict to sequences containing a complete open reading frame of at least 3,000 nucleotides. Examples in the prompt are selected by greedy farthest-point sampling of the 6-mer count vectors. The selected examples are concatenated with a ‘ATGCATGAATGCATGA’ separator. We evaluate 4, 8, 16, 24, and 32-shot prompts at temperature 0.9 and top_k 4.

We adopt the same correctness and novelty criteria from the first six filtering stages in Nguyen et al. (2024) to evaluate generations. These stages are: (1) an open reading frame of at least 3,000 nucleotides on either strand, (2) a Cas9 profile-HMM hit with *E* ≤ 10^−3^, (3) a CRISPR array detected by MinCED (Bland et al., 2007) of at least three repeats of 23 bases, (4) less than 90% amino acid identity to the natural loci using MMseqs2, using ‘-s 5.7 -e 0.001 –min-seq-id 0.0 -c 0.0 –cov-mode 0 –max-seqs 300 –split 0’, (5) a gap-length Gini coefficient ≤ 0.1 in a pairwise MAFFT (Katoh and Standley, 2013) alignment to the selected reference hit, (6) RuvC-I, RuvC-II, RuvC-III and HNH domain detected by HMM with *E* ≤ 10^−10^ within the gene.

We report the acceptance of the different filters. As a baseline we compare with the fine-tuned Evo 1 model published in Nguyen et al. (2024). When comparing pass-rates we pool all temperatures and prompt settings for both the fine-tuned and ICL generation. To compare the total number of hours to a library of 64 passing designs, we use the time of the previous total fine-tuning budget for the Evo 1 Cas9 fine-tune as well as estimated sampling time based on measuring generation using the Evo 1 codebase under the best temperature and setting, compared with the best ICL arm (Evo 2 7B ICL, 8 exemplars at *T* = 1.0, using the farthest point exemplar sampling).

### Mitochondria ICL

We collected pools of 32 mitochondrial genomes spanning different taxonomic groups (human, primate, mammal) from RefSeq (O’Leary et al., 2016). We selected only complete gnomes that were circular and contained 13 CDs and at least 24 RNAs. We sampled prompt sequences by greed farthest-point sampling using max-min cosine distance on 6-mer count vectors of the genomes. Full genomes are concatenated with a 24bp separator ‘ATGCATGAATGCATGAATGCATGA’ with no EOD token.

To evaluate generated sequences we use the vertebrate mitochondrial translation table 2 and call using six-frame stop segments of at least 30 amino acids using MMseqs2 against reference proteins from the downloaded mitochondria. tRNAs are annotated with tRNAscan-SE 2.0.13 (Chan et al., 2021) (‘-M vert’) and rRNAs using barrnap 0.9 (Seemann, n.d.) (‘–kingdom mito’). We evaluate if all 37 protein and RNA genes are present. To characterize novelty we measure the *k*-mer containment for 12-mers, which is what percent of the distinct 12-mers occur in the natural mitochondrial genome compared to the closest of all natural genomes in the pool. We also characterize the per-gene amino acid sequence divergence as the percentage difference to the nearest protein homology.

We compare the 2, 4, and 8-shot prompts on human and primate pools with Evo 2 20B, and then compare Evo 2 7B and Evo 2 20B on 8-shot prompts. We draw 10 samples per prompt with 4 randomized prompts per setting, using temperature of 0.7, top_k = 4, and top_p = 1, and generate a maximum of 20,000 bases or until the separator is emitted.

### Antibody ICL

We evaluate antibody design ICL on seven antigens: CNOT11 (PDB 8BFJ, chain A), IL-7RA (PDB 3DI3, chain A), PD-L1 (PDB 5O45, chain A), PVRIG (PDB 8X6B, chain A), SARS-CoV-2 RBD (PDB 6M0J, chain A), TNF (PDB 1TNF, chains A/B/C homo-trimer), and TRKA (PDB 1WWW, chain X). We use Germinal (Mille-Fragoso et al., 2026) to design scFVs against these targets with pre-specified hotspot positions, sampling until we collect 36 hits by the Germinal-established acceptance criteria (no AlphaFold 3 metrics are used to select hits to mitigate oracle leakage). All designs have 195 framework positions and 47 designable CDR positions at fixed indices (242 total). 8 seeds are selected from the hit pool for the prompt by greedy farthest-point selection on CDR identity. Each seed is reverse-translated by sampling codons from a human codon usage distribution. The 8 seeds are concatenated with a SEP token as the spacer to form the prompt.

ICL generation uses Evo2-flash with top-*k* 4, top-*p* 0.9, and temperatures of 0.9, 1.0, and 1.1 across 3 independent runs. Each generation is translated in the prompt reading frame, truncated to the first 242 residues, and designed CDRs are grafted onto the fixed framework. A generation is valid if it (1) reaches 242 residues, (2) carries no stop codons in CDR regions, and (3) has framework identity of at least 0.70 to the template before grafting.

We compare ICL to three baselines at a fixed budget of 1 GPU-hour for design and 1 GPU-hour for scoring. AbMPNN (Dreyer et al., 2023) is run through ColabDesign (Ovchinnikov et al., n.d.) (JAX implementation) to redesign the 8 seed backbones at the CDR regions, with all framework positions held fixed and the antigen present in complex during design. We sample with temperatures 0.1, 0.2, and 0.3 across 3 separate runs and omit cysteine, like the Germinal pipeline does during its AbMPNN step. The PSSM baseline samples each of the CDR positions independently from a position-specific distribution built from the 8 seeds. To make this baseline more realistic, we smoothed the distribution with a small symmetric Dirichlet prior, adding a pseudocount of 0.05 to every amino acid before normalizing, and again omitted cystine from the sampling alphabet. Twelve-thousand designs were drawn per antigen per temperature, since generation for this method is CPU-only and is effectively free. As a positive control for continuing to run the expensive generative model, we ran Germinal against the same targets with sequence design by AbMPNN at temperature 0.1.

For each generative method, designs are filtered and then subsampled to the folding budget. A design is retained if (1) its maximum CDR identity to the prompt union is at most 0.85, (2) its CDR H3 identity to the nearest prompt sequence is at most 0.70, and (3) it falls below 0.85 CDR identity to every design already kept. The order of designs is shuffled under a fixed seed before both steps so that the greedy scan used to compute filter step (3) is not biased.

For scoring, complexes were predicted with AlphaFold 3 (Abramson et al., 2024), as five diffusion samples per folding call. The binder is folded as a single sequence with no MSA, and the target chain uses an MSA. Of the five diffusion samples, the one with the highest AlphaFold 3 ranking score is kept, and every metric is computed from that sample. A design is considered a hit if it passes the following criteria: (1) mean pLDDT ≥ 0.70, (2) ipTM ≥ 0.50, (3) mean intra-binder pAE ≤ 13.0 Å, (4) pDockQ2 (Zhu et al., 2023) ≥ 0.15, (5) ipSAE (Dunbrack, 2025) ≥ 0.25, (6) fraction of interface residues that are CDR residues ≥ 0.40, (7) at least 3 CDR contacts, and (8) at least 1 hotspot contact.

To compute structural novelty, we use CDR RMSD where we superpose the framework alpha-carbons and compute the RMSD on the 47 CDR alpha-carbons. The threshold for novelty and distinct solution clustering is the median CDR RMSD across designs in our original Germinal-sampled pools.

To understand how ICL uses the prompt, each design’s 47 CDR positions were assigned to the seed each residue could have come from. Seed provenance is then assigned by a dynamic programming approach over (position, source) that minimizes the number of seed source switches.

### Bacteriophage genome ICL and CoT

We adopt the *Microviridae* dataset as well as correctness and novelty criteria on King et al. (2026). The filters are: (1) valid DNA sequence (A/C/G/T only) (2) length of 4,000-6,000 bases (3) GC content of 30-65% (4) no homopolymer longer than 10 nucleotides, (5) at least 7 genes with significant MMseqs2 (Steinegger and Söding, 2017) hits in the PHROG database (Terzian et al., 2021), and (6) 60-100% amino acid identity to the ΦX174 gene G. The novelty filters are then evaluated each separately for (1) a mean amino-acid identity of ≤ 95% across the best PHROG hits, (2) the genome architecture score, which measures the distance of start and stop of codons positions relative to natural ΦX174 and (3) the synteny criteria which requires the total and syntenic gene counts to be between 10-12, but excluding 11,11. These three novelty filters were also applied all together to show how higher diversity allows significantly higher pass rate for increasingly strict filters. The original implementations were adopted from King et al. (2026) for consistency, and the genomes generated by the fine-tuned Evo 2 checkpoint were used as a baseline, pooled across different hyperparameters.

For in-context learning we select from the 153 natural *Microviridae* genomes that pass the computational correctness filters. ICL prompts are selected with random selection among these 153 genomes; this is referred to as ICL on natural genomes in the results and figures. We sweep different numbers of genomes used in context, with 2, 4, 8 and 16, using temperature 0.7 and top_k = 4, adopting ‘ATGCATGAATGCATGA’ as the spacer to separate genomes and a maximum generation of 9,000 tokens if the spacer is not emitted.

For chain-of-thought generation, we start with appending generated genomes to the prompt in a first-in first-out manner with four genomes used in context. In the separate composition experiment we analyze the effect of different numbers of synthetic genomes in context for the *k* = 4 case, comparing the effects of having 0,1,2,3,4 synthetic genomes in context based on the cosine distance using normalized 6-mer count vectors measured to the closest prompt sequence and to the closest natural genome.

To construct the whole-genome phylogeny tree, we use IQ-Tree (Minh et al., 2020) on a whole-genome DNA alignment constructed by pyabpoa (Gao et al., 2021). All genomes are rotated to the circular origin and checked for compatible order and strand, with ΦX174 serving as the initialization.

To construct the t-SNE (Maaten and Hinton, 2008) visualization, we perform an all by all pairwise global Needlemean-Wunsch (Needleman and Wunsch, 1970) using Parasail (Daily, 2016) (gap-opening penalty 10, gap-extension penalty 1), with identity calculated by matching bases divided by alignment length, and t-SNE distances were 100 minus the percent sequence identity.

Capsid pentamers were predicted with AlphaFold 3, as five identical copies of each major-capsid call in a single job. Predictions were scored against the experimental ΦX174 capsid pentamer, generated by applying PDB 2BPA’s icosahedral assembly operators with Biotite (Kunzmann and Hamacher, 2018) and splitting the five symmetry copies into separate chains. Model-to-reference comparison used USalign in oligomeric mode (-mm 1 -ter 0), reporting TM-score normalized by the reference, RMSD and aligned length; interface confidence was taken from AlphaFold 3’s ipTM, with pTM, per-chain-pair ipTM, mean PAE and mean pLDDT (the B-factor column) recorded alongside (Abramson et al., 2024; Zhang et al., 2022; Zhang and Skolnick, 2004). Sequence identity to ΦX174 capsid protein F was computed by global optimal pairwise alignment (Biotite align_optimal, standard protein substitution matrix, affine gaps −10/−1).

To design MS2 and M13 phage, we develop correctness filters for these phage architectures. We adapt the gene calling method form King et al. (2026) to these phages, and create a pipeline based on (1) DNA sequence correctness (2) length within 20% of the natural genomes (MS2: NC_001417.2; M13: NC_003287.2), (3) at least one significant coding sequence as a MMseqs hit against the PHROG database, (4) recovery of every reference coding gene as a distinct, complete open reading frame. Complete ORFs were enumerated on both strands, and origin-spanning ORFs on the circular M13 genomes are retained, as well as overlapping or nested ORFs. Each annotated protein was required to have a length within ±20% of its reference protein, at least 30% amino-acid identity, and at least 90% alignment coverage of both the reference and candidate proteins.

To construct the prompts, we downloaded MS2 and M13 genomes from NCBI Virus and use the filters above to find only two MS2 genomes and six M13 genomes which pass our filters. Each prompt contained four M13 genomes or two MS2 genomes. For MS2 this means the same two genomes were used in every draw, although their order was shuffled. Complete example sequences were concatenated in the prompt using ‘ATGCATGAATGCATGA’. We generate sequences using Evo 2 7B with temperature 0.7 and top_k = 4, with maximum generation lengths of 10,000 tokens for M13 and 6,000 tokens for MS2. We generated eight sequences per prompt set across 126 prompt sets, yielding 1,008 sequences for each arm.

## Acknowledgements

We thank Brian Plosky for helpful discussions and assistance with manuscript preparation. We thank Brian Kang, Owen Tuck, Aayush Karan, Matthew Liu, and members of the Laboratory of Evolutionary Design for helpful discussions.

## Funding

G.B., A.T.M., and S.H.K. acknowledge funding support from the National Science Foundation Graduate Research Fellowship Program. A.T.M. acknowledges funding support from the Knight-Hennessy Graduate Scholarship Fund. B.L.H. acknowledges funding support for this project from Stanford Human-Centered Artificial Intelligence (HAI) Hoffman-Yee Research Grants, as well as broader research support from the Gates Foundation, the NIH Director’s New Innovator Award, the Advanced Research Projects Agency for Health (ARPA-H), Arc Institute, Schmidt Sciences AI2050, Stanford Center for Digital Health, and Silicon Valley Community Foundation. Lanmodulin design work was supported by the U.S. Department of Energy (DOE), Office of Science, Office of Biological and Environmental Research project award number SCW1929. Contributions from LLNL were performed under the auspices of the U.S. Department of Energy by Lawrence Livermore National Laboratory under contract DE-AC52-07NA27344 (LLNL-JRNL-2026228).

## Author Contributions

G.B., Y.A.H., and B.L.H. conceived the study. B.L.H. supervised the project. G.B. and Y.A.H. implemented and tested the Evo 2 vLLM engine. G.B. conducted the CoT study, and designed and analyzed Cas9 and phage generations with inputs from S.H.K. Y.A.H. designed and analyzed LanM and Rubisco, with inputs from K.E.S. and D.M.P. D.N. designed and analyzed antibodies with inputs from T.W. A.T.M. designed and analyzed Acrs.

## Competing Interests

B.L.H. acknowledges outside interest in Arpelos Biosciences and Genyro Inc. as a scientific co-founder. No other competing interests are declared.

## Data, Code, and Materials Availability

Code and model checkpoints for vLLM-accelerated Evo 2 (Evo2-flash) are available at https://github.com/ ArcInstitute/evo2 and https://github.com/AlexYkHao/evo2-vllm.

## Supplementary Figures

**Fig. S1.**
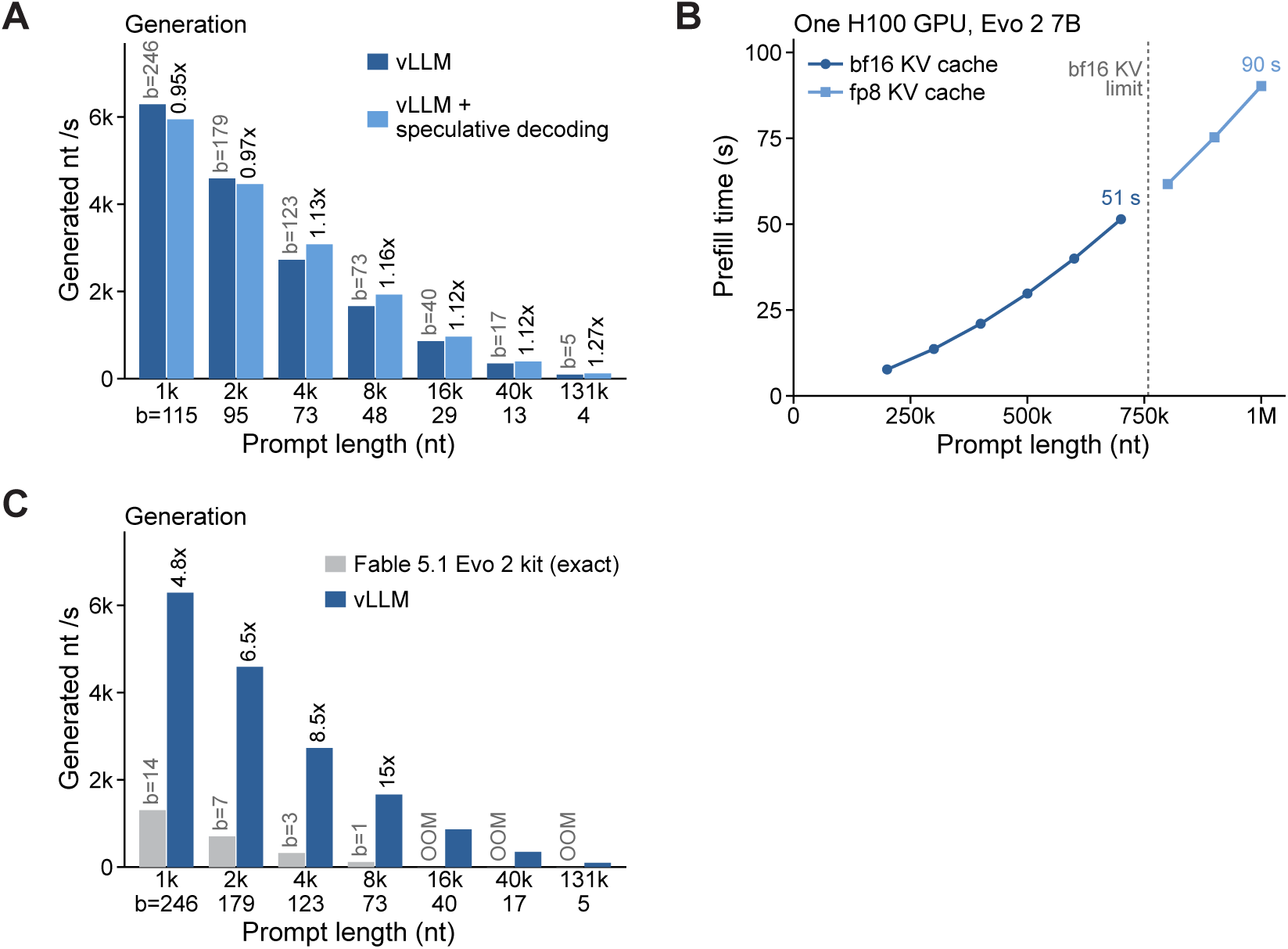
Evo2-flash is faster and more memory-efficient. **(A)** The trained speculative drafter further speeds up the Evo2-flash inference engine in the long prompt regime (prompt longer than 4 kilobases). For decoding, we used the largest batch size allowed by the GPU. The variable *b* indicates the batch size used in each condition. **(B)** Evo2-flash can fit up to 1 megabase on one H100 (80GB) GPU, with fp8 KV cache. With bf16 KV cache it fits up to 700 kilobases. **(C)** Evo2-flash speeds up generation and relaxes the memory constraint compared with the Evo 2 kit reported in the Claude Fable 5.1 system card (Anthropic, 2026). For decoding, we used the largest batch size allowed by the GPU.

**Fig. S2.**
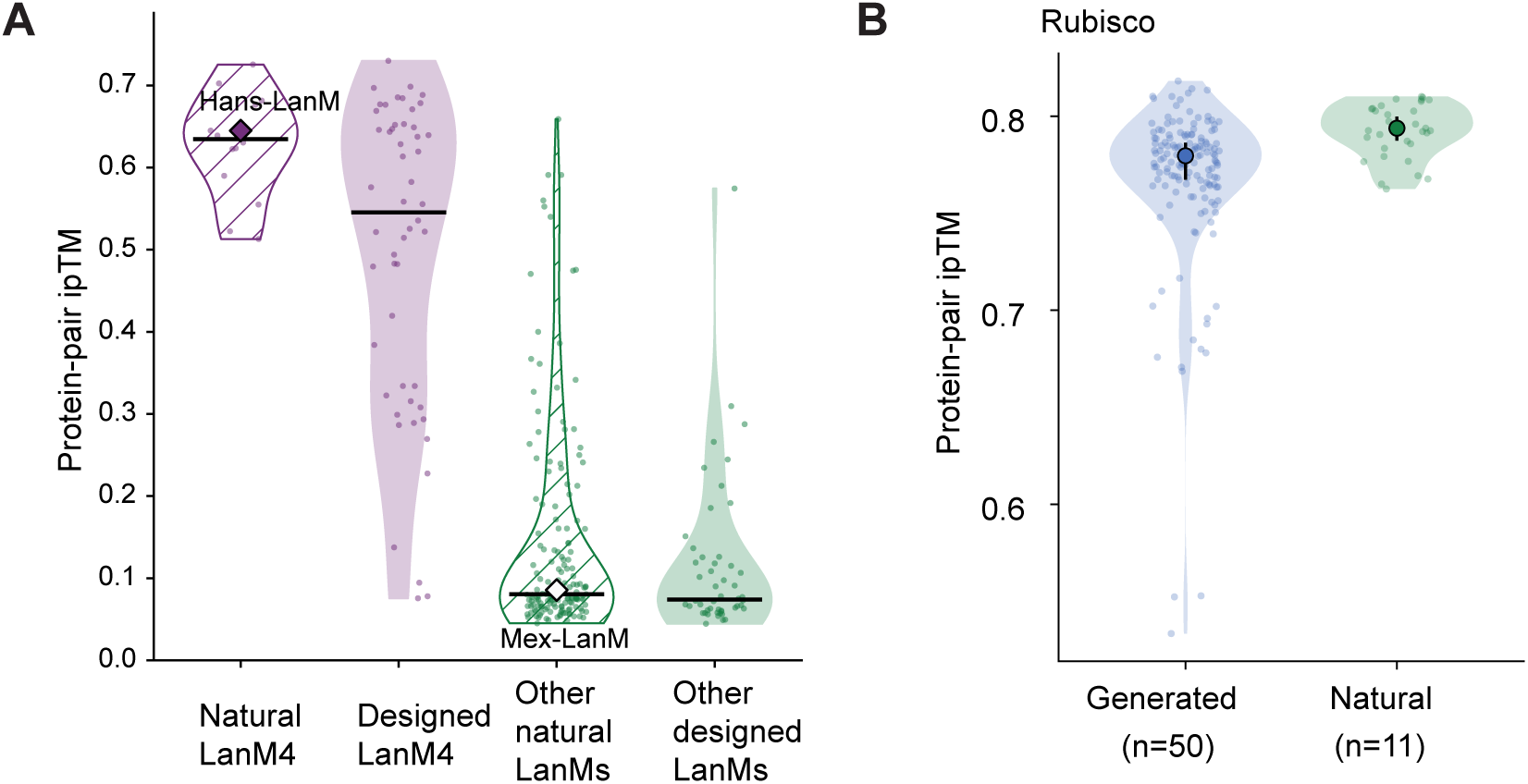
Predicted intermolecular interactions of natural and designed proteins. **(A)** ESMFold2-modeled intermolecular interactions of natural and designed LanMs. For reference, Hans-LanM was reported to be homodimeric by experiments, and Mex-LanM was reported to be monomeric (Mattocks et al., 2023). We filtered out sequences with pLDDT less than 75. **(B)** ESMFold2-modeled intermolecular interactions of natural and designed rubiscos. We filtered out sequences with pLDDT less than 75.

**Fig. S3.**
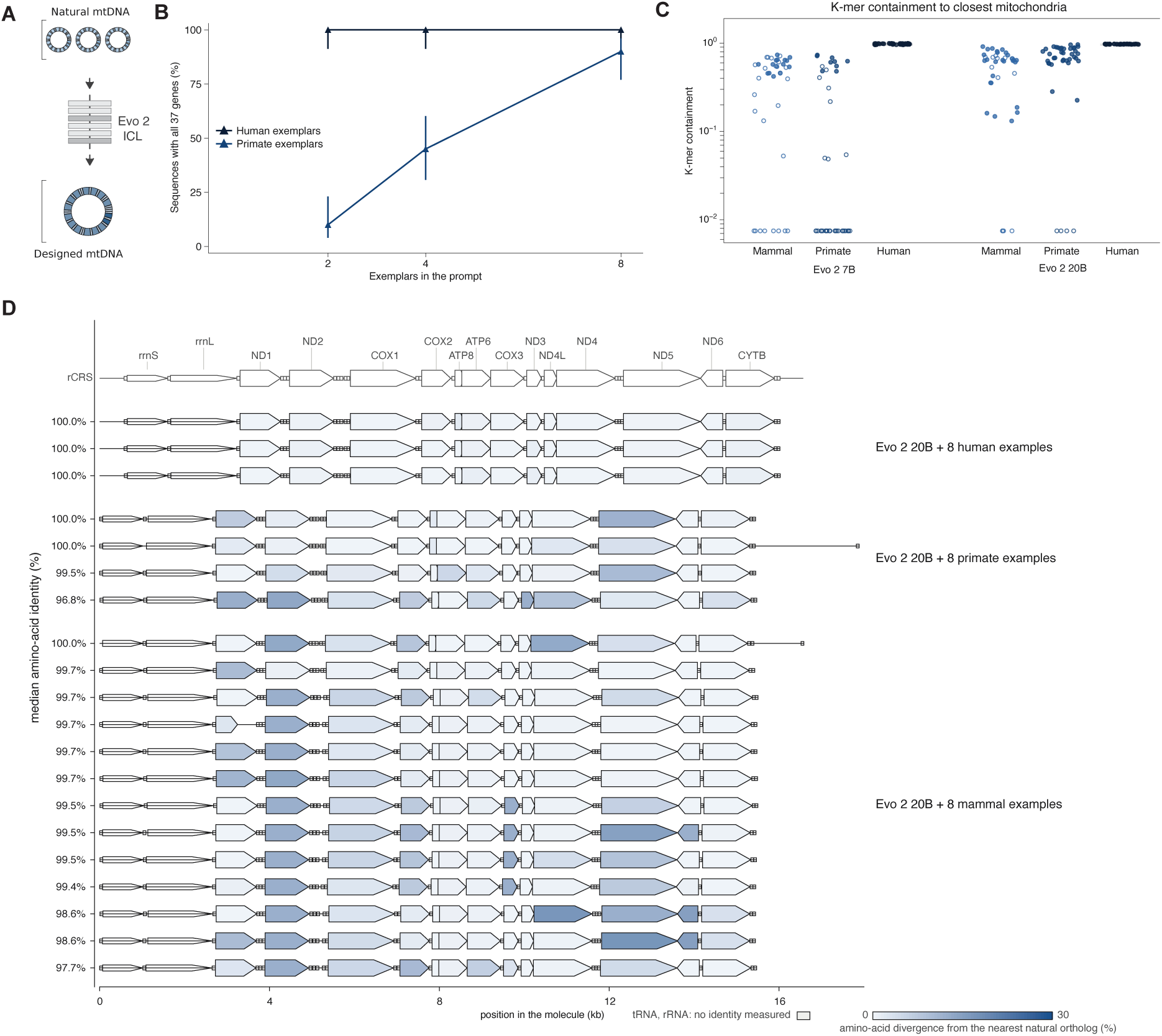
Mitochondria ICL genome designs. **(A)** Schematic of mitochondrial genome design using ICL. **(B)** Percentage of generated sequences containing all 37 mitochondrial genes as a function of the number of exemplars in the prompt, comparing human and primate exemplars. **(C)** The *k*-mer containment (*k* = 12) of the generated mitochondrial genomes compared with the closest natural genome in the dataset. **(D)** Gene maps of mitochondrial genomes generated by Evo 2 20B with 8 human, primate, or mammal examples, shown against the human reference (rCRS). Protein-coding genes are colored by amino acid divergence from the nearest natural ortholog; tRNAs and rRNAs are not colored. Values at left indicate the median amino acid identity of each genome.

**Fig. S4.**
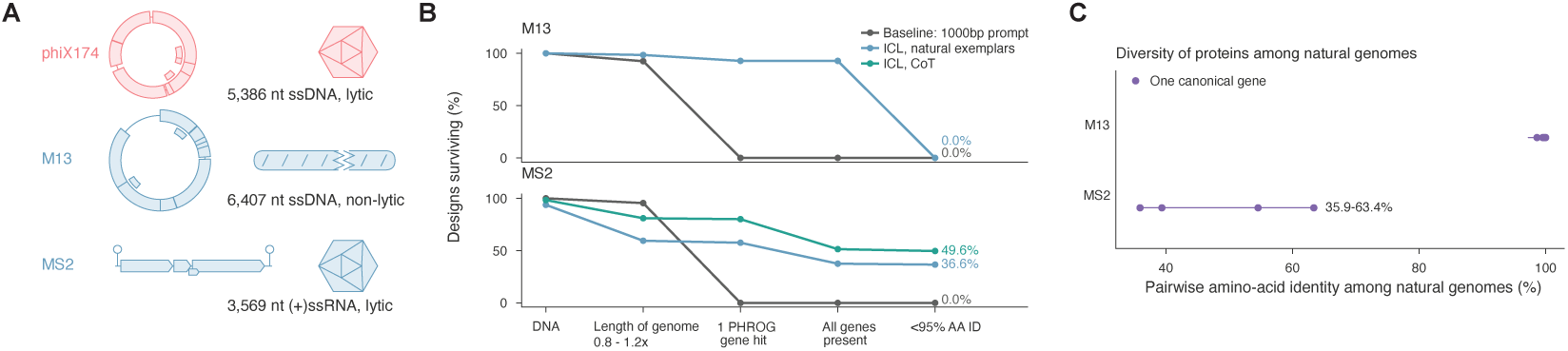
MS2 and M13 phage ICL genome designs. **(A)** Schematic showing the genome architecture, capsid schematic, and lifestyles of the M13 and MS2 phages compared with ΦX174. **(B)** Percentage of designs passing each filtering stage using Evo 2 7B ICL and CoT compared with prompting Evo 2 with a 1,000 bp prompt. **(C)** The natural diversity of proteins between different M13 and MS2 genomes by pairwise amino acid identity of the same gene.

